# Precise Control of Sodium Alginate Microdroplets through pH-Sensitive Cross-Linkers and Geometric Factors

**DOI:** 10.1101/2025.09.08.674978

**Authors:** Alireza Rezvani, Mehrnaz Oveysi, Mohammad Mahdi Karim khani, Vahid Bazargan, Marco Marengo

## Abstract

This study presents a novel approach to achieve internal and on-chip gelation of sodium alginate microdroplets using pH-sensitive water-insoluble cross-linkers. Gelation is initiated by precisely controlling the introduction of acetic acid, which decreases the pH, through the generation of microdroplets via step emulsification. Unlike current approaches in microfluidic droplet generation, where droplet size depends on various external factors such as flow rate ratio and fluid properties, our strategy utilizes the device’s geometry to regulate droplet size. Importantly, this approach ensures that there is no aggregation in the channels following microgel creation. The empirical investigation focuses on exploring the impact of geometrical factors on alginate droplet formation in our customized microfluidic device. To expedite the production of polydimethylsiloxane (PDMS) microfluidic devices, we utilized 3D-printed molds, which provide effective control over channel height. Using our method, we successfully generate microdroplets with a coefficient of variation (CV) below 3% and diameters ranging from 100 to 200 µm. It is worth noting that these microdroplets experience an average shrinkage of 16% following gelation, indicating the successful formation of stable gels. By combining pH-sensitive cross-linkers, precise control over acetic acid introduction, and the geometric regulation of droplet size, our approach demonstrates great potential for the development of microfluidic devices capable of producing uniform and stable microdroplets for various applications, including drug delivery, tissue engineering, and biotechnology.

## Introduction

In recent years, the advancement of microfluidic technologies has revolutionized various fields, offering unprecedented precision and efficiency in manipulating small volumes of fluids [1]. Among these innovations, integrating three-dimensional (3D) printing techniques into microfluidic device fabrication has emerged as a promising approach, eliminating traditional lithography methods and providing control, particularly in the z-direction [2]. In this paper, we explore the utilization of 3D-printed resin molds for fabricating polydimethylsiloxane (PDMS)-based microfluidic devices for droplet generation. By leveraging the capabilities of 3D printing to produce intricate resin molds, we have optimized the microfluidic device design to facilitate precise droplet generation. A recent study by Anyaduba et al. [3] used the same protocol for mold fabrication, further validating the effectiveness and applicability of this approach in microfluidic device manufacturing. Specifically, we have capitalized on the inherent advantages of the resin mold’s 3D-printed structure to incorporate features such as height variations within the microfluidic channels [4]. The study conducted by Wu et al. exemplifies the incorporation of structures with varying heights in a microfluidic system. They employed a combination of micro-stereolithography 3D printing and CNC micro-milling to create microwells with depths ranging from 100 to 500 µm.

Our droplet generation model, termed “step emulsification” [6], operates on the spontaneous transformation mechanism. Using this methodology, we have attained exceptional uniformity in droplet size and enhanced control over the droplet generation process. Step emulsification offers robustness against variations in flow rate ratios, ensuring consistent droplet sizes irrespective of flow rate fluctuations [7]. In step-emulsification microfluidic devices, the droplet neck pinch-off process occurs without the requirement of shearing and squeezing the continuous phase. However, it is essential to have enough supply of the continuous phase to ensure the timely removal of droplets near the orifice [8].

Alginate microgels have displayed intriguing characteristics across various fields and applications, spanning from medicinal to environmental domains [9], [10]. Utilizing droplet-based techniques for their fabrication provides enhanced control over their production. We employed the microdevice to produce alginate droplets, capitalizing on the inherent differences in microdevice heights to achieve consistent droplet sizes. The cross-linking of these alginate droplets can be achieved through different methods, either internally or externally. Internal crosslinking involves adding insoluble or poorly soluble cross-linking agent salts to the alginate solution and inducing gelation with acidified oils. Alternatively, external crosslinking entails adding the cross-linking agent to pre-formed alginate microdroplets [11]. In our approach, we introduced calcium carbonate nanoparticles into the alginate solution to internally initiate gelation of the droplets [12].

### Device Fabrication

The master molds’ 3D models were created using SolidWorks CAD software (see Figure 1), These models were subsequently printed using the Water-Washbale Rapid Black 3D Printing resin (Phrozen), with a layer thickness of 25 µm. Following printing, the master molds underwent cleaning with isopropyl alcohol and were finally subjected to UV-curing for 45 minutes. The PDMS precursor (SYLGARD® 184) from Sigma Aldrich, was blended with a curing agent in a 10:1 (w/w) ratio and subjected to degassing in a desiccator for over 30 minutes. Following this, the PDMS prepolymer mixture was poured onto the resin mold and cured in an oven at 75°C for more than 2 hours. Once solidified, the PDMS replica was carefully removed from the resin mold. Inlets and outlets were then created by punching holes using a tool with a diameter of 1.5 mm. Finally, the PDMS replica was sealed with a glass slide (76 mm × 26 mm; thickness 0.9–1.2 mm) through oxygen plasma treatment (power: 20 W; pressure: 1 Torr) for 15 s.

**Figure 1.**
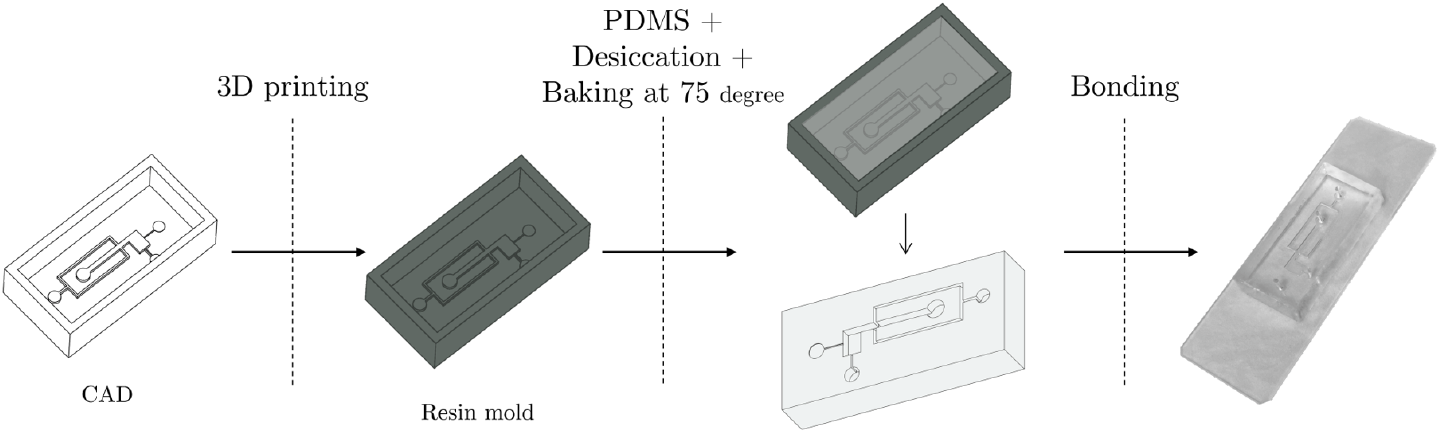
Microchip Manufacturing Process Overview: Steps include 3D modeling of master molds using SolidWorks, SLA 3D printing, PDMS casting and curing, inlet/outlet creation, and sealing with a glass slide via oxygen plasma treatment.

## Material

Light mineral oil (kinematic viscosity at 25°C = 20.2−22.2 cSt), Sodium Alginate (Product number W201502, kinematic viscosity in water at 25°C = 19.4 cSt), Acetic acid, and surfactant (Span 80) were procured from Sigma-Aldrich Chemicals, USA. The Calcium Carbonate nanopowder was sourced from Nanosany Corporation.

### Configuration of the microfluidic device

As depicted in Figure 2, the step height, denoted as *h*^∗^, ranged from 3 to 20 µm in our experiments for different molds. The illustration of the mold design is shown, with the inlet channel height and width set at 100 µm.

**Figure 2.**
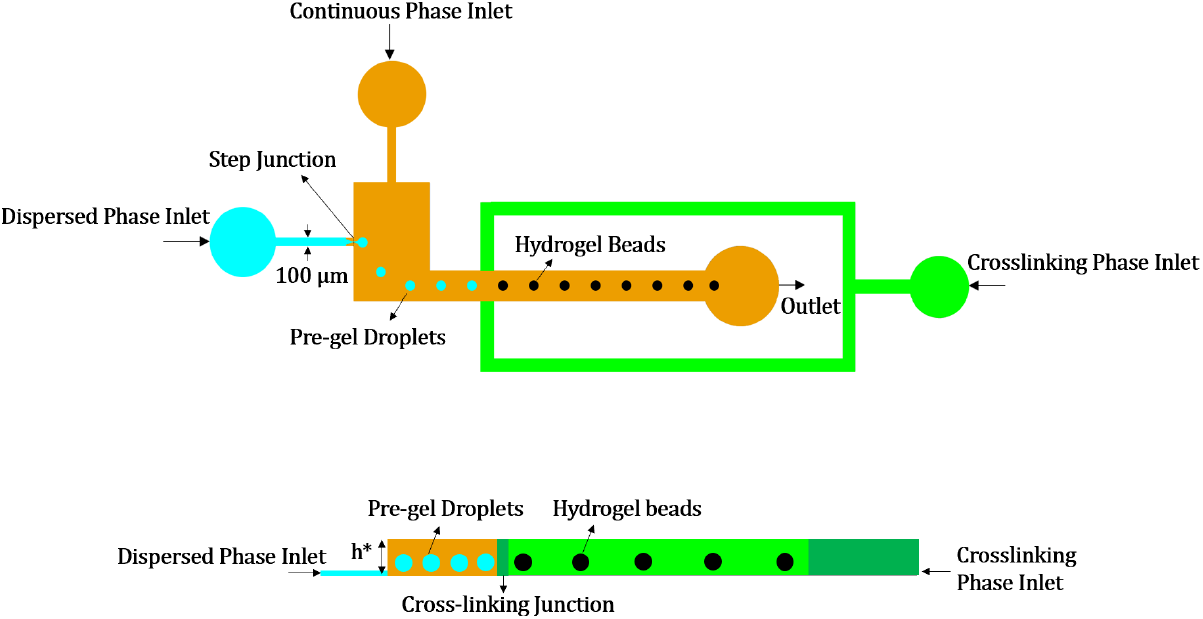
Microfluidic Chip setup Depicting Injection of Phases Leading to Gel Formation: Top view displaying the injection points of different phases culminating in gel formation; Side view inset showcasing the height differential within the chip

Before proceeding to examine the gelation mechanism and to characterize droplet formation in our microfluidic device, we introduced the two phases separately into the microfluidic chip using a syringe pump (Harvard PHD ULTRA) set at the specified flow rate. The dispersed phase (*Qd*) consisted of DI water or an aqueous alginate solution, while the continuous phase (*Qc*) was composed of oil with 3wt% Span 80. (*Qc*) is fixed at 0.4 µL/min in the whole experiment. The visualization was captured using a high-speed Pixelink camera (Model Number: PL-D7912CU-T), which was mounted on an Olympus IMT-2 fluorescent inverted microscope. Direct microscopy images and the Pixelink Scope software were utilized to determine the size of droplets and microgels. To measure the sizes accurately, calibration with a known reference was performed. Subsequently, the area of the particles was measured using the circle tool provided by the software, and the corresponding diameter was calculated from the area measurement. Following iterative experiments and thorough characterization to define the optimal flow rate and microfluidic device design, we employed the established parameters to fabricate droplets. Gelation was initiated by injecting a mixture of mineral oil and acetic acid in the crosslinking phase inlet as depicted in the Figure 2.

## Results and discussion

### Flow regime diagram

In the step emulsification process, two distinct flow patterns for droplet formation have been identified in our experiments: the dripping regime, characterized by the production of monodisperse droplets at the step and the balloon droplet formation, wherein droplet pinch-off does not occur at the junction, instead exhibiting mass accumulation and droplet expansion. Subsequently, due to the flow of the continuous phase, droplet pinch-off occurs post-expansion, akin to observations in a T-junction configuration. These two regimes are depicted in Figure 3, showcasing the stages of droplet formation in each regime over time. Additionally, there are instances where droplet formation fails to occur, characterized by the expansion of the dispersed phase without subsequent break-up.

**Figure 3.**
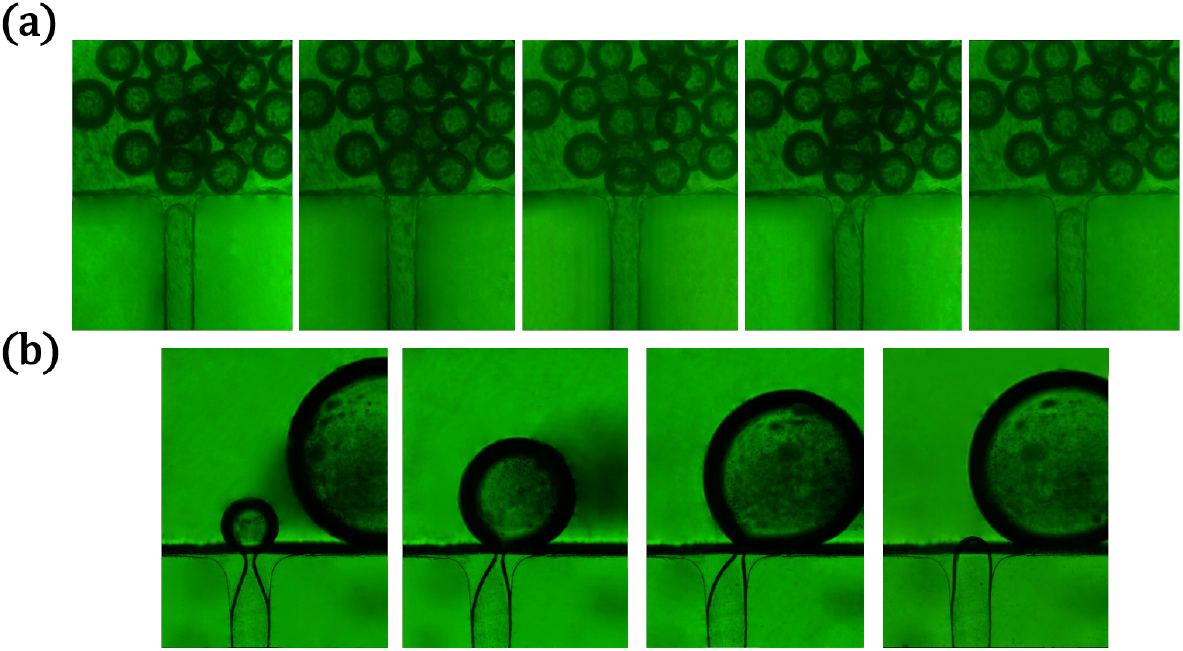
Illustration of Droplet Formation Regimes - (a) Dripping regime showcasing monodisperse droplet generation at the step; (b) Balloon droplet formation characterized by mass accumulation and subsequent droplet expansion

To characterize our microfluidic device regarding droplet formation regimes, we conducted two experiments using different dispersed phases: first, deionized (DI) water, followed by an aqueous solution of alginate (1wt%). Introducing a dimensionless parameter, *h*, defined as *h*^∗^ (the step height) divided by the inlet channel height, we plotted *h* against the flow rate of the dispersed phase. Each data point on this plot corresponds to a specific droplet formation regime. Figure 4 illustrates this plot, delineating the regions of dripping, balloon droplet formation, and no droplet formation for (a) DI water and (b) aqueous solution of alginate as the dispersed phase. Raising the step height in both cases expands the dripping region and postpones balloon formation. We observe that the dominant regime of droplet formation is dripping, even at high flow rates of the dispersed phase, when water is used as the dispersed phase. However, for alginate, the range of dripping is narrower, requiring a higher step height and lower flow rate of dispersed phase to ensure dripping. This can be attributed to the solution’s higher viscosity compared to water, which increases the viscous forces resisting droplet breakup.

**Figure 4.**
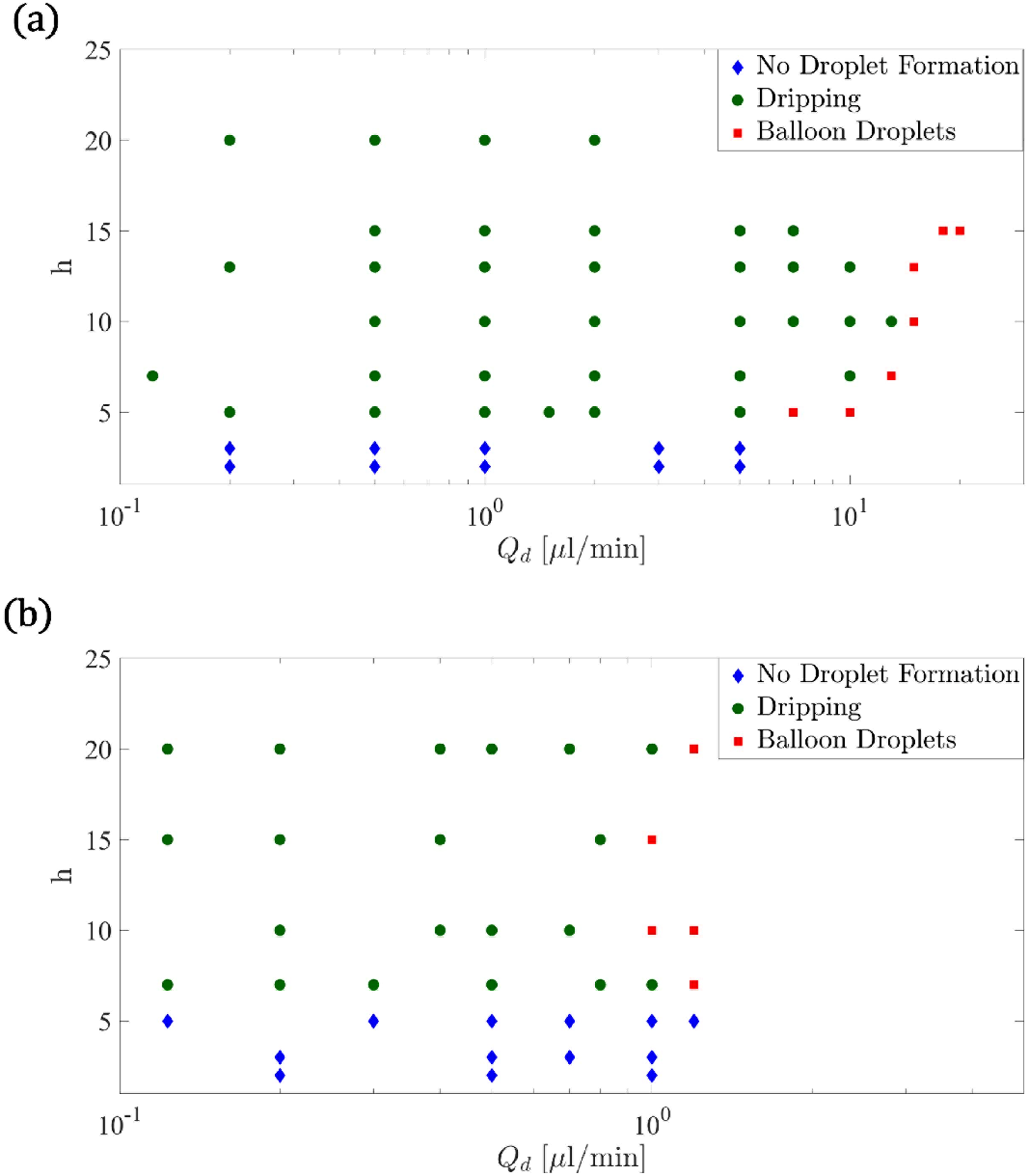
Flow Regime Diagram for Droplet Formation - Mapping dimensionless parameter *h* against dispersed phase flow rate for (a) DI water and (b) Aqueous solution of alginate. The plot delineates distinct regions representing dripping, balloon droplet formation, and no droplet formation.

### Droplet Size and Formation Frequency: Effects of Step Height and Dispersed Phase Viscosity

One of the crucial advantages of employing microfluidic droplet generators is their capability to produce droplets under controlled conditions, allowing for precise manipulation of droplet size and production rate. We investigated this aspect by plotting the diameter of water and alginate droplets against the flow rate of the dispersed phase. The experiments were conducted using microfluidic chips of varying heights. As depicted in Figure 5, increasing the step height or the flow rate of the dispersed phase results in the production of larger droplets. Specifically, when water was used as the dispersed phase, droplets with diameters exceeding 200 µm were produced, facilitated by the ability to employ higher flow rates. In contrast, alginate droplets typically ranged between 100 to 200 µm in diameter. It is worth noting that without any alteration to the continuous phase, droplet size could be controlled solely by adjusting the geometrical aspects of the microchip [13]. In contrast to other geometries such as the well-known flow-focusing method, where estimating droplet size requires correlation and scaling formulas accounting for factors like flow rate ratio of dispersed and continuous phases, capillary number of the continuous phase, and viscosity ratio [14], our introduced microdevice offers a simplified approach. Here, droplet size can be estimated solely as a function of the dispersed flow rate for each step height. This streamlined estimation process underscores the effectiveness and efficiency of our microdevice for droplet generation

**Figure 5.**
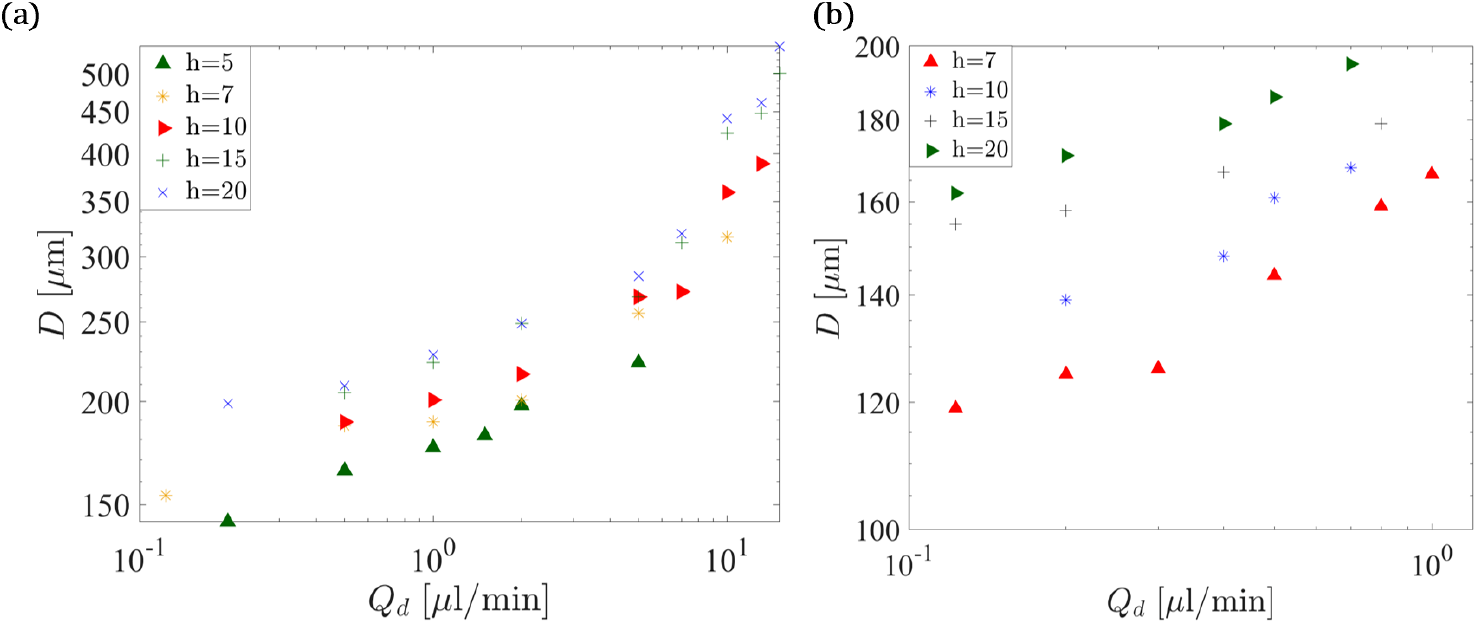
Influence of Step Height and Dispersed Phase Flow Rate on Droplet Size for (a) DI Water and (b) Alginate Aqueous Solution - Increasing step height or dispersed phase flow rate leads to larger droplet formation.

In Figure 6, we observe the variation in droplet generation frequency with dispersed phase flow rate for both DI water (a) and an aqueous solution of alginate (b). We note that the frequency initially increases with the flow rate, but as we approach the balloon regime, it begins to decrease. In the dripping regime, droplet separation at the step junction is primarily driven by surface tension and instability induced by height variation. However, as we transition towards the balloon regime, the momentum force of the internal phase intensifies, resulting in more forceful droplet detachment. Consequently, as droplet size increases significantly, the frequency of droplet production declines. Additionally, we observe that increasing the height of the step leads to a decrease in droplet production frequency. This is attributed to the heightened difficulty in cutting larger droplets as the step height increases, resulting in larger droplet diameters.

**Figure 6.**
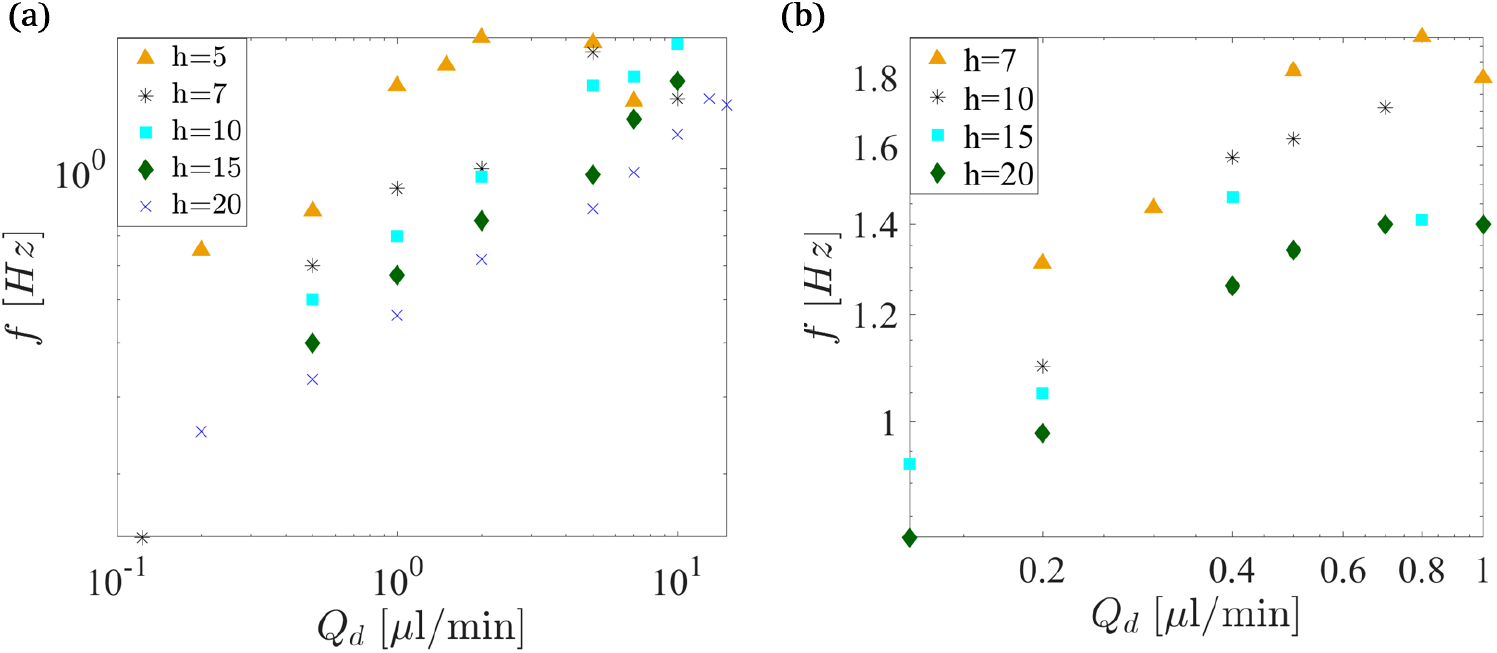
Variation in Droplet Generation Frequency with Dispersed Phase Flow Rate for(a) DI Water and (b) Alginate Aqueous Solution

### Alginate microgel formation

After characterizing our microdevice for alginate microdroplet formation and understanding the effects of h and dispersed phase flow rate on droplet size and production rate, we can select the optimal step height and flow rate of the dispersed phase to produce alginate microdroplets of the desired size. Subsequently, after injecting the mixture of acetic acid and alginate as described earlier, on-chip gelation occurred. The microgels can then be transferred to an aqueous solution of calcium chloride (2 wt%) to complete the gelation process. Figure 7 illustrates the on-chip droplets and the resulting microgels generated from them. The generated microdroplets shown in Figure 7 are highly monodispersed, with a coefficient of variation (CV) below 3%, calculated using the formula 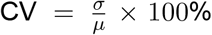, where *σ* is the standard deviation and *µ* is the mean diameter of the droplets. The microgel experienced a reduction in volume of 16% after complete gelation, determined using the shrinkage formula: Shrinkage 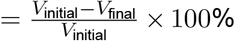, where *V*_initial_ represents the initial volume of the microgel and *V*_final_ denotes the volume of the microgel after gelation.

**Figure 7.**
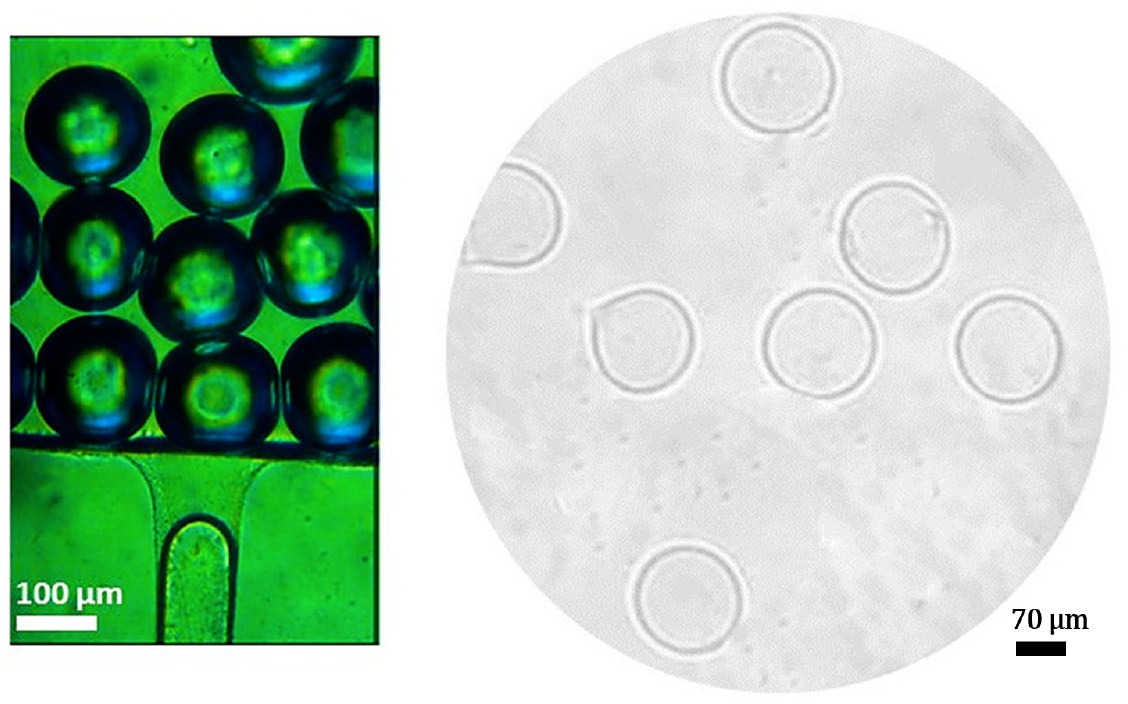
Microfluidic Generation of Monodisperse Alginate Droplets (178 µm) and Subsequent Gelation into Microgels (153 µm)-The step height was chosen as 20 µm and flow rate of dispersed phase was 0.4 µL/min.

As depicted in Figure 1, for alginate droplet formation, we selected a step height of 20 µm. In determining this height, we also considered the probability of droplet merging after formation with the previously fabricated droplet. As the step height increased, this effect diminished, resulting in a lower probability of merging. We hypothesize that the reduction in surface energy due to the formation of larger droplets contributes to the increased stability of the droplets with increasing height.

## Conclusions

1. Our study showcases the efficacy of microfluidic technology in precise control over droplet size and formation frequency, achieved through the manipulation of microchip geometrical aspects.
2. Both step height and flow rate of the dispersed phase are identified as critical parameters influencing droplet size, with larger droplets produced by increasing either of these factors. Leveraging our optimized conditions, we successfully fabricated highly monodisperse alginate microdroplets and observed minimal variation in size, with a coefficient of variation below 3%. Furthermore, we extended the functionality of our microfluidic device by transforming microdroplets into microgels directly on the chip. This integration of gelation processes within the microfluidic platform offers a streamlined and efficient approach to producing microgels with precise control over size and morphology,
3. Through a systematic approach in microchip manufacturing, including 3D modeling, SLA 3D printing, PDMS casting, and hermetic sealing, we have established a robust platform for future microfluidic applications.

### Nomenclature

Please use three-line chart shown above to display the nomenclatures and abbreviations.

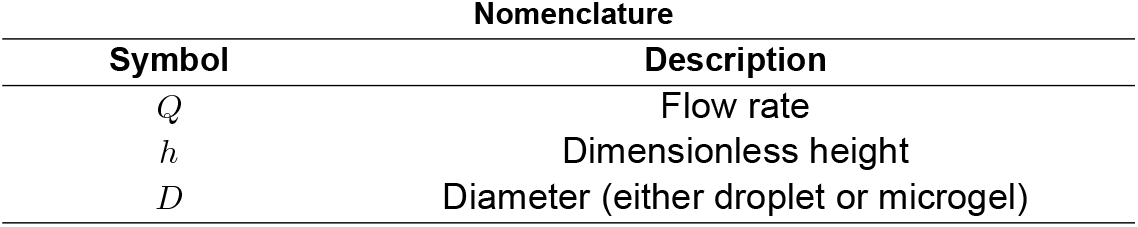

## References

[1] S. S. H. Tsai, Y. L. Lin, P. C. Huang, and J. Y. Wu. Recent advances in microfluidic technologies for biological applications: A review. Journal of Biomedical Science and Engineering, 2023.

[2] J. C. Miller and T. D. Nguyen. Three-dimensional printing of microfluidic device molds: Advances and challenges. Lab on a Chip, 2022.

[3] T. D. Anyaduba, J. A. Otoo, and T. S. Schlappi. Picoliter droplet generation and dense bead-in-droplet encapsulation via microfluidic devices fabricated via 3d printed molds. Micromachines, 13(11): 1946, 2022.

[4] Pingan Zhu and Liqiu Wang. Passive and active droplet generation with microfluidics: a review. Lab on a Chip, 17(1): 34–75, 2017.

[5] E. Behroodi, H. Latifi, Z. Bagheri, et al. A combined 3d printing/cnc micro-milling method to fabricate a large-scale microfluidic device with the small size 3d architectures: an application for tumor spheroid production. Scientific Reports, 10:22171, 2020.

[6] M. A. M. Abate and D. A. Weitz. Step emulsification. Lab on a Chip, 2011.

[7] Z. Liu, C. Duan, S. Jiang, C. Zhu, Y. Ma, and T. Fu. Microfluidic step emulsification techniques based on spontaneous transformation mechanism: A review. Journal of Industrial and Engineering Chemistry, 2020.

[8] Wei Zhan, Ziwei Liu, Shaokun Jiang, Chunying Zhu, Youguang Ma, and Taotao Fu. Comparison of formation of bubbles and droplets in step-emulsification microfluidic devices. Journal of Industrial and Engineering Chemistry, 106: 469–481, 2022.

[9] Amin Ghaffari, Alireza Rezvani, Taha Goudarzi, and Ehsan Amani. How does the interconnector design influence coronary stents structural and hemodynamic performance? Journal of the Brazilian Society of Mechanical Sciences and Engineering, 47(1): 28, 2025.

[10] Silvia Vicini et al. Alginate gelling process: Use of bivalent ions rich microspheres. Polymer Engineering & Science, 57(6): 531–536, 2017.

[11] C. B. Goy and R. E. Chaile. Microfluidics and hydrogel: A powerful combination. React. Funct. Polym., 145:104314, 2019.

[12] S. Akbari and T. Pirbodaghi. Microfluidic encapsulation of cells in alginate particles via an improved internal gelation approach. Microfluidics and Nanofluidics, 16: 773–777, 2014.

[13] Amin Ghaffari, Mobina Amiri Vahid, Pedram Zaree, Kasra Mazarei Saadabadi, and Mahsa Ghaffari. Blood flow analysis of subject-specific cerebral arterial tree: A focus on the redistribution of the blood flow after occlusion. In 2025 IEEE 22nd International Symposium on Biomedical Imaging (ISBI), pages 1–5. IEEE, 2025.

[14] J Tan, JH Xu, SW Li, and GS Luo. Drop dispenser in a cross-junction microfluidic device: Scaling and mechanism of break-up. Chemical Engineering Journal, 136(2–3): 306–311, 2008.

